# Development of a competition assay to assess the *in vitro* fitness of dengue virus serotypes using an optimized serotype-specific qRT-PCR

**DOI:** 10.1101/2024.09.10.611934

**Authors:** Anne-Fleur Griffon, Loeïza Rault, Etienne Simon-Lorière, Myrielle Dupont-Rouzeyrol, Catherine Inizan

## Abstract

**Background:** Comparing the *in vitro* fitness of dengue virus (DENV) isolates is a pivotal approach to assess the contribution of DENV strains’ replicative fitness to epidemiological contexts, including serotype replacements. Competition assays are the gold standard to compare the *in vitro* replicative fitness of viral strains. Implementing competition assays between DENV serotypes requires an experimental setup and an appropriate read-out to quantify the viral progeny of strains belonging to different serotypes.

**Results:** In the current study, we optimized an existing serotyping qRT-PCR by adapting primer/probe design and multiplexing the serotype-specific qRT-PCR reactions, allowing to accurately detect and quantify all four DENV serotypes. The qRT-PCR was specific, had a limit of detection of at least 5.08×10^1^, 5.16×10^1^, 7.14×10^1^ and 1.36 x10^1^ genome copies/µL, an efficiency of 1.993, 1.975, 1.902, 1.898 and a linearity (R²) of 0.99975, 0.99975, 0.9985, 0.99965 for DENV-1, -2, -3 and -4 respectively. Challenge of this multiplex serotype-specific qRT-PCR on mixes of viral supernatants containing known concentrations of strains from two serotypes evidenced an accurate quantification of the amount of genome copies of each serotype. We next developed an *in vitro* assay to compare the replicative fitness of two DENV serotypes in the human hepatic cell line HuH7: quantification of the viral progeny of each serotype in the inoculum and the supernatant using the serotype-specific multiplex qRT-PCR unveiled an enrichment of the supernatant in DENV-1 genome copies, uncovering the enhanced replicative fitness of this DENV-1 isolate.

**Conclusions:** This optimized qRT-PCR combined to a relevant cellular model allowed to accurately quantify the viral progeny of two DENV strains belonging to two different serotypes in a competition assay, allowing to determine which strain had a replicative advantage. This reliable experimental setup is adaptable to the comparative study of the replicative fitness of any DENV serotypes.

## Background

Dengue viruses are RNA viruses classified into four serotypes (DENV-1 to -4) based on their antigenic properties. DENV serotypes share between 34 and 73% identity at the nucleotide sequence level ^1^. The emergence of a new serotype has the potential to lead to outbreaks of enhanced breadth and severity, some DENV serotypes, genotypes or clades being associated with different severity, especially upon secondary infections ^2^. However, the mechanisms underlying DENV serotypes co-circulation and replacements are misunderstood. Viral replicative fitness may contribute to DENV serotype replacements. It is therefore of prime importance to set up experimental models to assess the relative replicative fitness of strains from different serotypes. For RNA viruses, an approximation to fitness is the relative ability to produce stable infectious progeny in a given environment ^3^. The study of viral replicative fitness has become an expanding field of research ^4^. DENV fitness was initially approached through *in vitro* monoinfections of cellular models and quantification of the viral progeny or yield of virus, either by qRT-PCR or titration ^5,6^.

For many other viruses, the established gold standard to compare the replicative fitness is the setting up of a competition assay, which allowed to compare the fitness of two variants of various viruses, including HIV-1 and influenza viruses ^4^, Saint Louis Encephalitis virus ^7^, Bovine Viral Diarrhea Virus 1 and 2 ^8^ or mutants of Mayaro virus ^9^ for instance. The two viral variants or types were specifically quantified in the supernatant using either titration in the presence of variant-specific monoclonal antibodies ^7^, a flow cytometry assay using strain-specific fluorescent probes ^8^ or variant/type-specific primers and probe ^7-9^. A search of PubMed using the following query ((dengue virus) AND ((competition) OR (coinfection)) AND ((cell) OR (cellular) OR (in vitro))) returned 259 results, among which many refer to competition and super-infection assays in cellular models between DENV and other arboviruses, including yellow fever virus ^10^, chikungunya virus ^11^, Sindbis Virus ^12^, Zika virus ^13^ or Japanese encephalitis virus and insect-specific viruses ^14^. These assays were probed either with virus-specific qRT-PCR ^11-13^ or flow-cytometry using virus-specific antibodies ^14^. Competition in mosquito C6/36 cells between two DENV serotypes were also reported; they were probed using type-specific hyper-immune mouse ascites ^15,16^, monoclonal antibodies ^16^ or a reverse transcription followed by a SybR Green qPCR using serotype-specific primers ^17^. Finally, competition between two DENV NS4B mutants was reported and quantified by measurement of peak height at the polymorphic position in Sanger sequencing along with 3D-digital PCR with mutant-specific primers and probes ^18^.

Quantification by molecular techniques appears as the most time- and cost-efficient strategy to quantify the progeny yielded by strains belonging to two different serotypes. Several serotype-specific qRT-PCR have been developed to serotype DENV, especially for diagnostic purposes ^19,20^. The challenge is to leverage these techniques to allow the absolute quantification of the viral progeny of a cell culture simultaneously infected by strains belonging to two different serotypes. Furthermore, most competition assays implemented with DENV were developed in mosquito cells, recapitulating only part of the viral life cycle. In the current study, we aim to develop an *in vitro* competition assay in human cells to compare the relative replicative fitness of two DENV strains belonging to two different serotypes using a serotype-specific qRT-PCR read-out.

## Methods

### Cell culture

The *Aedes albopictus* C6/36 cell line was cultured at 28°C in Leibovitz L15 medium (Sigma-Aldrich, Merck, Steinheim, Germany) supplemented with 10% decomplemented Fetal Calf Serum (FCS, Gibco™, Fisher scientific, Paisley, UK) and 10% Tryptose Phosphate Broth (TPB, Gibco™, Fisher scientific, Paisley, UK). The human hepatocarcinoma HuH7 cell line was cultured at 37°C under 5% CO_2_ in Dulbecco’s Modified Eagle Medium (DMEM, Gibco™, Fisher scientific, Paisley, UK) supplemented with 10% decomplemented FCS.

### Production of DENV viral isolates

DENV isolates were recovered from DENV positive human serum samples through infection of C6/36 cells. All final viral stocks were prepared with no more than four passages in C6/36 cells for 5 days. Supernatants were collected and stored at −80 °C in 0.5 M Sucrose (Acros Organics, Geel, Belgium) and 20 mM Hepes buffer (Gibco, Fisher scientific, Paisley, UK). Viral isolates were titrated by Immunofluorescent Focus Assay as described previously ^21^; their titers were expressed in Focus Forming Unit per mL (FFU/mL). Viral isolates were recovered for two DENV-1 genotype I, two DENV-2 genotype Cosmopolitan, 7 DENV-3 genotype I and one DENV-4 genotype IIa. Characteristics of the viral isolates are given in Table 1.

**Table 1.** Characteristics of the dengue viral isolates used in this study.

| **Serotype** | **Name of the strain** | **Year of collection** | **Concentration (genome copies/μL)** | **Titer (FFU/mL)** | **Used in monoplex qRT-PCR** | **Used for color compensation** | **Used in multiplex qRT-PCR** | **Used for standard curve** | **Used in mixes** | **Used in *in vitro* infections** |
| --- | --- | --- | --- | --- | --- | --- | --- | --- | --- | --- |
| DENV-1 | AVS73<sup>a</sup> | 2017 |  |  | X | X |  |  |  |  |
|  | AVS94<sup>b</sup> | 2017 | 6.03x10<sup>5</sup> | 3.93x10<sup>7</sup> |  |  | X | X | X | X |
| DENV-2 | AVS109<sup>c</sup> | 2017 |  |  | X | X |  |  |  |  |
|  | DWL089<sup>c</sup> | 2018 | 3.74x10<sup>5</sup> | 1.24x10<sup>7</sup> |  |  | X | X | X | X |
| DENV-3 | DWL133<sup>c</sup> | 1995 |  |  | X |  |  |  |  |  |
|  | 409 | 2008 |  |  | X |  |  |  |  |  |
|  | 240<sup>c</sup> | 2014 |  |  | X | X |  |  |  |  |
|  | 5734 | 2014 |  |  | X |  |  |  |  |  |
|  | 5519 | 2014 |  |  | X |  |  |  |  |  |
|  | DWL039<sup>c</sup> | 2017 | 4.94x10<sup>5</sup> | 1.67x10<sup>7</sup> | X |  | X | X | X | X |
|  | DWL049<sup>c</sup> | 2017 |  |  | X |  |  |  |  |  |
| DENV-4 | 8626 | 2009 |  |  | X | X |  |  |  |  |

<sup>a</sup> GenBank accession number MW315187, <sup>b</sup> GenBank accession number MW315194, <sup>c</sup> GISAID EPI\_SET\_240904sd doi:
[10.55876/gis8.240904sd](https://doi.org/10.55876/gis8.240904sd)

### RNA extraction

RNA extraction from serum samples was performed using the QIAamp Viral RNA extraction kit (Qiagen, Hilden, Germany) following the manufacturer instruction. Viral RNA were stored at -80°C until subsequent use.

### Quantification of the RNA concentration of DENV viral isolates by qRT-PCR

The absolute concentrations of DENV viral isolates were measured by Taqman qRT-PCR using the primers and the probe described in ^19^. The reaction mix contained 0.3 µM of primers, 0.07 µM of FAM/BHQ-1-coupled probe, 1.2 µL of Superscript III Platinum One-Step enzyme (ThermoFisher Scientific, Carlsbad, CA, USA), 15 µL of the corresponding buffer (Invitrogen) and 5 µL of viral RNA extract in a reaction volume of 30 µL. Thermal cycling protocol consisted in a reverse transcription step of 20 min at 50°C, followed by an enzyme activation and initial denaturation step of 2 min at 95°C, after which 50 cycles of 5 s of denaturation at 95°C and 30 s of annealing and extension at 60°C were implemented. Fluorescence was collected at the end of each annealing and extension step in the 465/510 nm canal. qRT-PCR reactions were performed using a LightCycler 480 (Roche, Basel, Switzerland). Absolute quantification of the number of copies genome per µL were performed using an RNA calibrator (TibMolBiol, Berlin, Germany). The calibrator sequence is given in the supplementary materials.

A standard curve was established by performing 10-fold dilutions of the RNA calibrator in water, from 10^8^ to 10^2^ copies/µL. Each dilution of the RNA calibrator was loaded in duplicate in the PCR mix described above and run using the thermal cycling protocol described above. The resulting standard curve was analyzed using the LightCycler 480 software (Supplementary figure 1). In each subsequent qRT-PCR, a sample containing 10^5^ genome copies/µL of the RNA calibrator (median concentration of RNA calibrator detected in the standard curve) was loaded and run in parallel of the RNA extracts to quantify. Concentrations were calculated using the standard curve previously established and the LightCycler 480 software.

### Optimization of a serotype-specific multiplex qRT-PCR

In order to quantify the absolute concentrations of DENV viral isolates belonging to two different serotypes in a mixture of two viral strains, we adapted and optimized a serotype-specific singleplex Taqman qRT-PCR ^20^. Initial ^20^ and optimized primers and probes (Eurogentec, Seraing, Belgique) are detailed in Table 2.

**Table 2.**
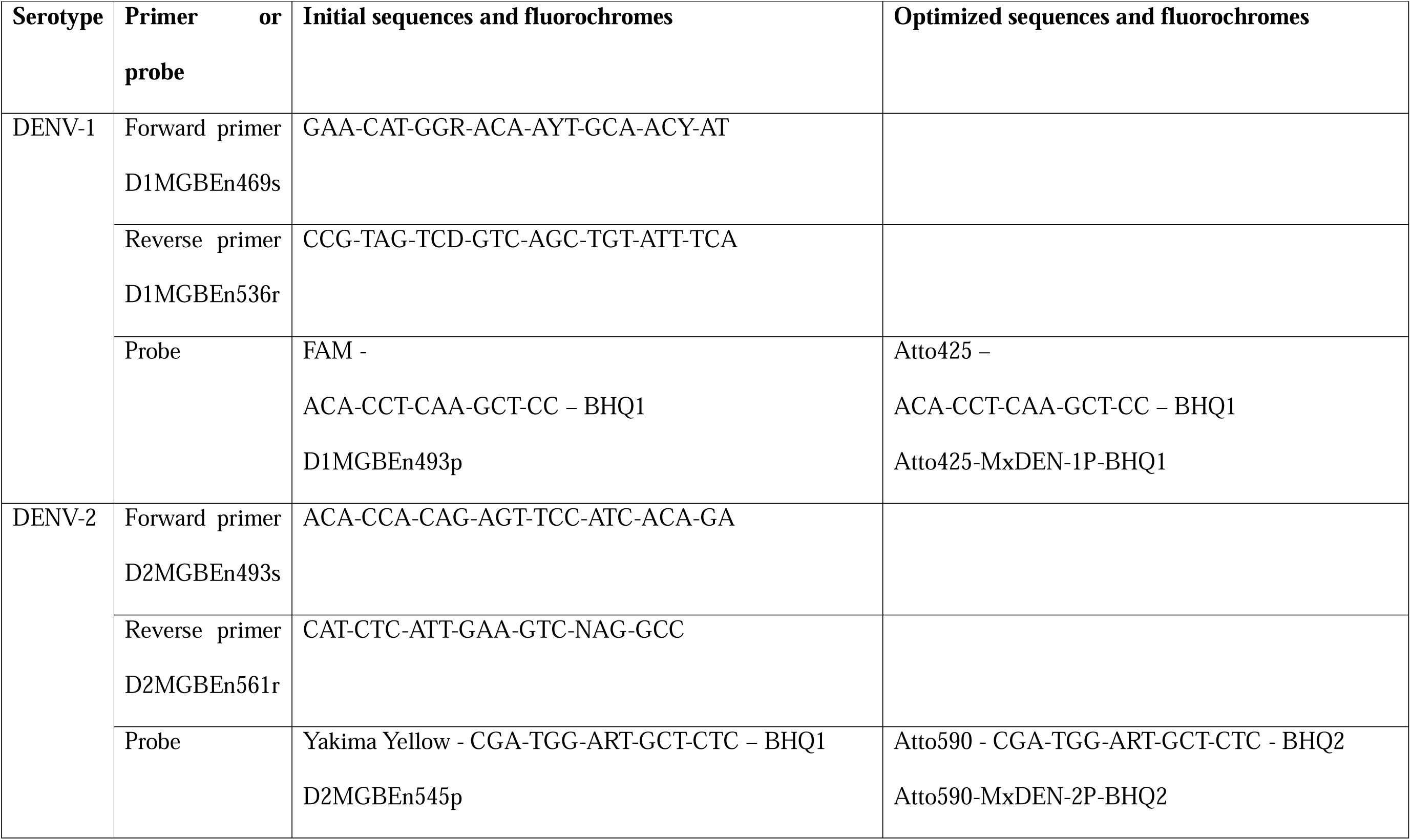
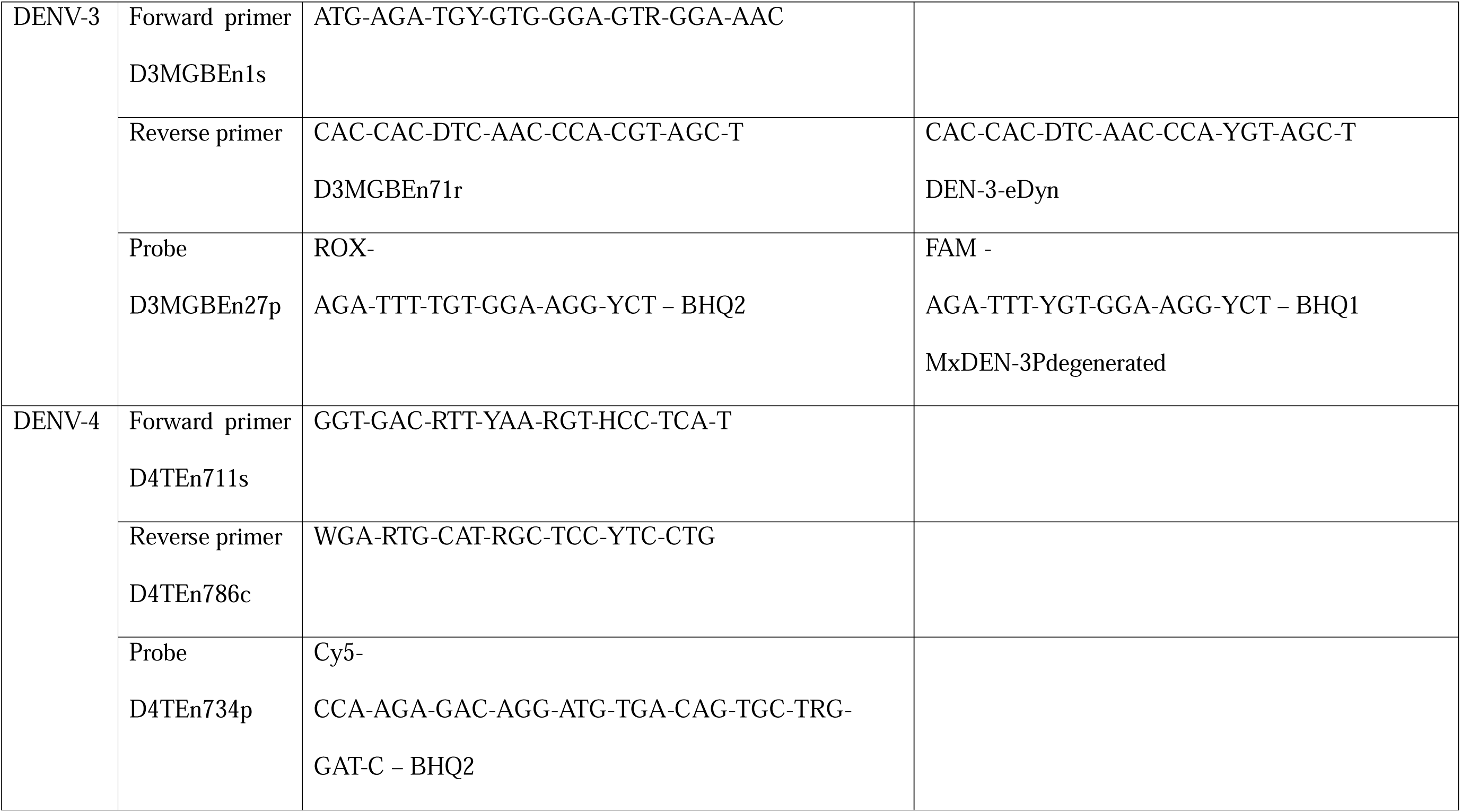
Sequences of the primers and probes and fluorochrome combinations.

The composition of multiplex reaction mixes in primers and probes, allowing the detection and quantification of all 4 DENV serotypes, is described in Table 3. Additionally, reaction mixes contained 0.8 µL of Superscript III Platinum One-Step enzyme (Invitrogen), 10 µL of the corresponding buffer (ThermoFisher Scientific, Carlsbad, CA, USA) and 4 µL of RNA extract in a reaction volume of 20 µL.

**Table 3.** Final concentrations of primers and probes in the optimized serotype-specific qRT-PCR.

| **Serotype** | **Primer or probe** | **Final concentration**<br>**(μM)** |
| --- | --- | --- |
| DENV-1 | Forward primer D1MGBEn469s | 0.28 |
|  | Reverse primer D1MGBEn536r | 0.28 |
|  | Probe D1MGBEn493pa | 0.125 |
| DENV-2 | Forward primer D2MGBEn493s | 0.28 |
|  | Reverse primer D2MGBEn561r | 0.28 |
|  | Probe D2MGBEn545pa | 0.50 |
| DENV-3 | Forward primer D3MGBEn1s | 0.28 |
|  | Reverse primer D3MGBEn71r | 0.28 |
|  | Probe D3MGBEn27pa | 0.50 |
| DENV-4 | Forward primer D4TEEn711s | 0.28 |
|  | Reverse primer D4TEEn786c | 0.28 |
|  | Probe D4TEEn734pb | 0.50 |

Thermal cycling protocol consisted in a reverse transcription step of 20 min at 50°C, followed by an enzyme activation and initial denaturation step of 2 min at 95°C, after which 50 cycles of 5 s of denaturation at 95°C and 30 s of annealing and extension at 60 °C were implemented. Fluorescence was collected at the end of each annealing and extension step in the 440/488, 498/580, 533/610 and 618/660 nm canals for DENV-1, DENV-3, DENV-2 and DENV-4, respectively. qRT-PCR reactions were performed using a LightCycler 480 (Roche, Basel, Switzerland). A color-compensation experiment was performed on the Light Cycler 480 according to the manufacturer instructions. Briefly, for each of all 4 DENV serotypes, the RNA extract was loaded in pentaplicates in the corresponding qRT-PCR reaction mix containing all 8 primers and the probe specific to this serotype. The thermal cycling protocol consisted in a reverse transcription step of 20 min at 50°C, followed by an enzyme activation and initial denaturation step of 2 min at 95°C, after which 50 cycles of 5 s of denaturation at 95°C and 30 s of annealing and extension at 60 °C were implemented. The samples were then placed 1s at 95°C with a ramping of 4.4°C/s, then 30s at 40°C with a ramping of 2.2°C/s and underwent a ramping of 0.14°C/s up to 65°C during which a continuous acquisition of the fluorescence at each increase in 1°C was conducted in the 440/488, 498/580, 533/610 and 618/660 nm canals. Viral isolates of the four serotypes used for this color-compensation experiment are specified in Table 1. Following color compensation calculations, fluorescence detection thresholds of qRT-PCR results were manually adjusted based on the signal yielded by the serotype-specific positive controls (Supplementary figure 2).

### Absolute quantification of the RNA concentrations of DENV viral isolates using the serotype-specific multiplex qRT-PCR

Viral isolates of the four serotypes used for this quantification are specified in Table 1. The absolute concentration of these RNA extracts was determined by qRT-PCR as described in a previous section. A standard curve was established by performing 10-fold dilutions of these RNA extracts in water. Each dilution of the RNA extract was loaded in duplicate in the PCR mix described and the thermal cycling protocol described above. The resulting standard curve was analyzed using the LightCycler 480 software (Supplementary figure 3).

Absolute quantification of the number of genome copies per µL was then performed using these RNA extracts of known concentrations as calibrators: in each subsequent qRT-PCR, a sample of each of these in-house calibrators containing 5.08×10^3^, 5.16×10^3^, 7.14×10^3^ and 1.36×10^3^ genome copies/µL for DENV-1, DENV-2, DENV-3 and DENV-4, resp., was loaded and run in parallel of the RNA extracts to quantify. Concentrations were calculated in the Light Cycler 480 software using the corresponding standard curve previously established.

### Preparation of mixes of viral isolates from two different serotypes at a determined ratio

Viral isolates of known concentration were artificially mixed two by two to generate viral mixes containing 100:0, 75:25, 50:50, 25:75 and 0:100 ratios in genome copies of two DENV strains belonging to two distinct serotypes. Viral isolates used are specified in Table 1. Viral RNA was extracted from these artificial viral mixes as described in a previous section and concentrations in viral RNA from each serotype were quantified by serotype-specific multiplex qRT-PCR.

### Competition assay

HuH7 were seeded at 2 x 10^5^ cells per well in 12-well plates. Twenty-four hours later, cells were infected with DENV viral isolates diluted in DMEM supplemented with 2% FCS to a final volume of 300 µL. Cells were infected either in “mono-infections” by a single viral isolate or in “competition” by a mix of two viral isolates. Multiplicities of infection (MOI) were calculated based on the titer of the viral isolate in FFU/mL determined by Immunofluorescent Focus Assay. For mono-infections, cells were infected with a single viral isolate at a MOI of one. For competitions, cells were infected with a mix of two isolates belonging to two different serotypes at a MOI of one for each isolate (ratio 50:50) or at a MOI of 0.2 for one isolate and 1.8 for the other isolate (ratio 10:90). After two hours of incubation at 37°C under 5% CO_2_, the inoculum was retrieved and stored at -80°C in 0.5 M Sucrose and 20 mM Hepes buffer for subsequent RNA extraction. Cells were washed twice with 1 mL of DMEM supplemented with 2% FCS. Cells were then cultured at 37°C under 5% CO_2_ in 1 mL of DMEM supplemented with 2% FCS. At 72h post-infection, supernatants were collected and stored at -80°C in 0.5 M Sucrose and 20 mM Hepes buffer for subsequent RNA extraction.

## Results

### qRT-PCR optimization to detect all four DENV serotypes by qRT-PCR

Primers and probes of a serotype-specific qRT-PCR^20^ were coupled with a combination of fluorochromes compatible with multiplexing on the LC480 (Table 2): DENV-1 probe was coupled to FAM, DENV-2 probe to Yakima Yellow, DENV-3 probe to ROX and DENV-4 probe to Cy5 (Table 2). Positions of primers and probes on DENV genome are given in Figure 1A. In first intention, they were challenged in singleplex qRT-PCR for the accurate detection of DENV-1 to DENV-4 strains retrieved in New Caledonia between 1995 and 2018 (Table 1). DENV-1, -2 and -4 strains tested were accurately detected (Supplementary figure 4A, B and D). DENV-3 retrieved in 1995, 2008 and 2014 were also detected, albeit with a low fluorescence signal (Supplementary figure 4C). DENV-3 strains retrieved in 2017 were not detected (Supplementary figure 4C).

**Figure 1.**
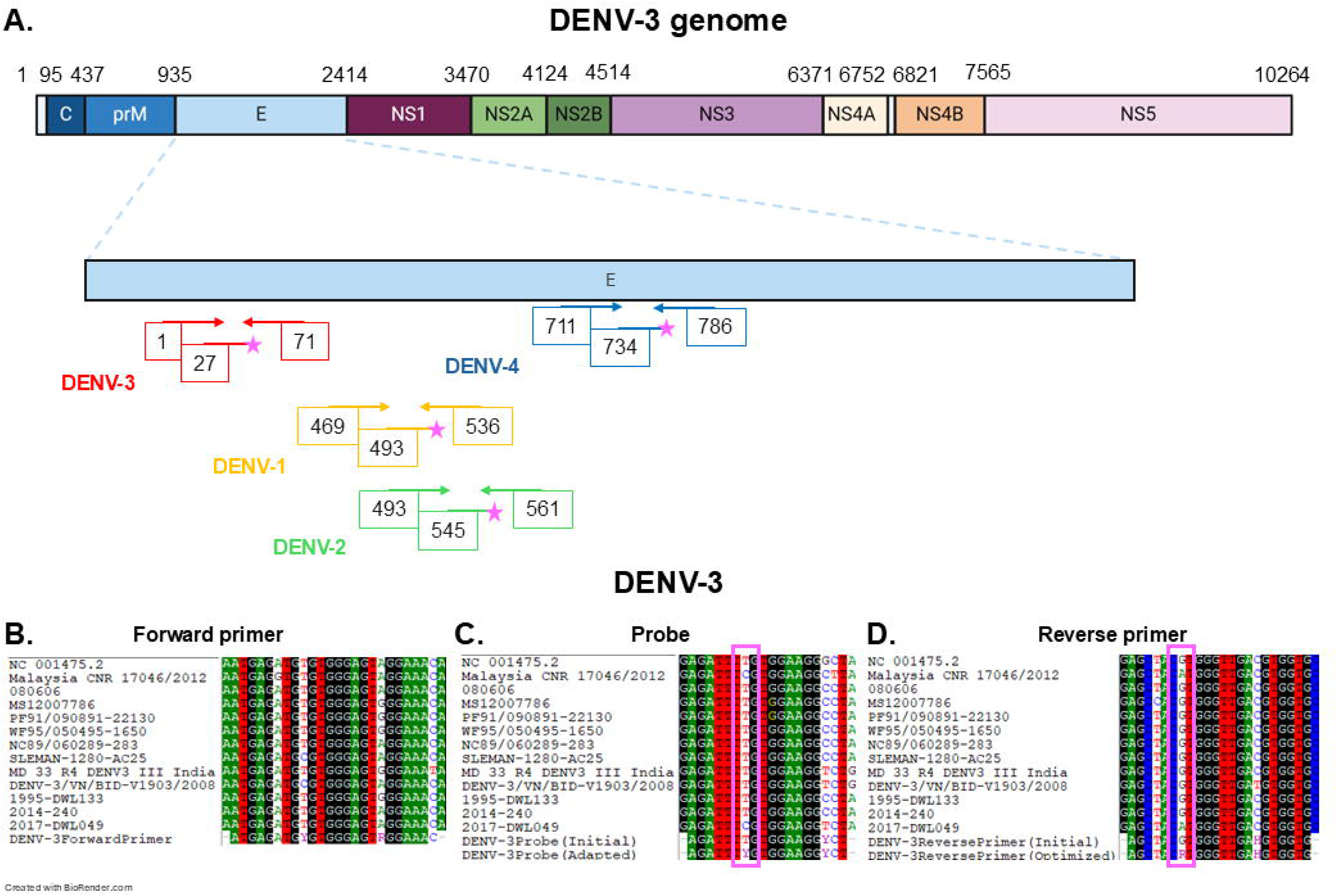
Design of optimized primers and probes. **A.** Positions of the 5’ end of DENV-1 to -4 primers and probes on DENV-3 genome, as indicated in Table 2. Positions are given according to DENV-3 genome annotation (strain NC_001475.2). Pink stars represent the fluorochromes on the probes. **B-D.** Alignments of DENV-3 forward primer (**B.**), initial and adapted probe (**C.**) and initial and adapted reverse primer (**D.**) with DENV-3 sequences from genotypes I, II and III, including DENV-3 genotype I strains DWL133, 2014-240 and 2017-DWL049 retrieved in New Caledonia in 1995, 2014 and 2017 respectively. Optimized positions are highlighted with a pink frame.

Alignment of DENV-3 primers and probes with DENV-3 sequences from genotype I, II and III including DENV-3 genotype I strains retrieved in 2017 in New Caledonia evidenced a perfect match for the forward primer (Figure 1B). However, this alignment revealed two mismatches with DENV-3 strains retrieved in NC in 2017 and in Malaysia: a mismatch at position 7 in the probe (C in DENV-3 VS T in the probe, Figure 1C) and a mismatch at position 16 in the reverse primer (A in the DENV-3 VS G in the reverse complemented sequence of the reverse primer, Figure 1D). Sequences of the probe and reverse primer were optimized to include degenerated nucleotides able to detect the diversity of DENV-3 strains at these positions: a Y was introduced in the probe sequence at position 7, and a Y was introduced in the reverse primer sequence at position 16. Furthermore, the combination of fluorochromes was optimized for detection on the LightCycler 480 by coupling DENV-1 probe to Atto425, DENV-2 probe to Atto590, DENV-3 probe to FAM and DENV-4 probe to Cy5 (Table 2). Optimized primers and probes were challenged in singleplex qRT-PCR for the accurate detection of DENV-1 to DENV-4 strains retrieved in New Caledonia between 1995 and 2018 (Table 1). All DENV-1 to -4 strains tested were accurately detected with a strong fluorescence signal (Figure 2A.-D.). Only DENV-2 fluorochrome Atto590 yielded a maximal fluorescence intensity of 0.64 AU (Figure 2B.).

**Figure 2.**
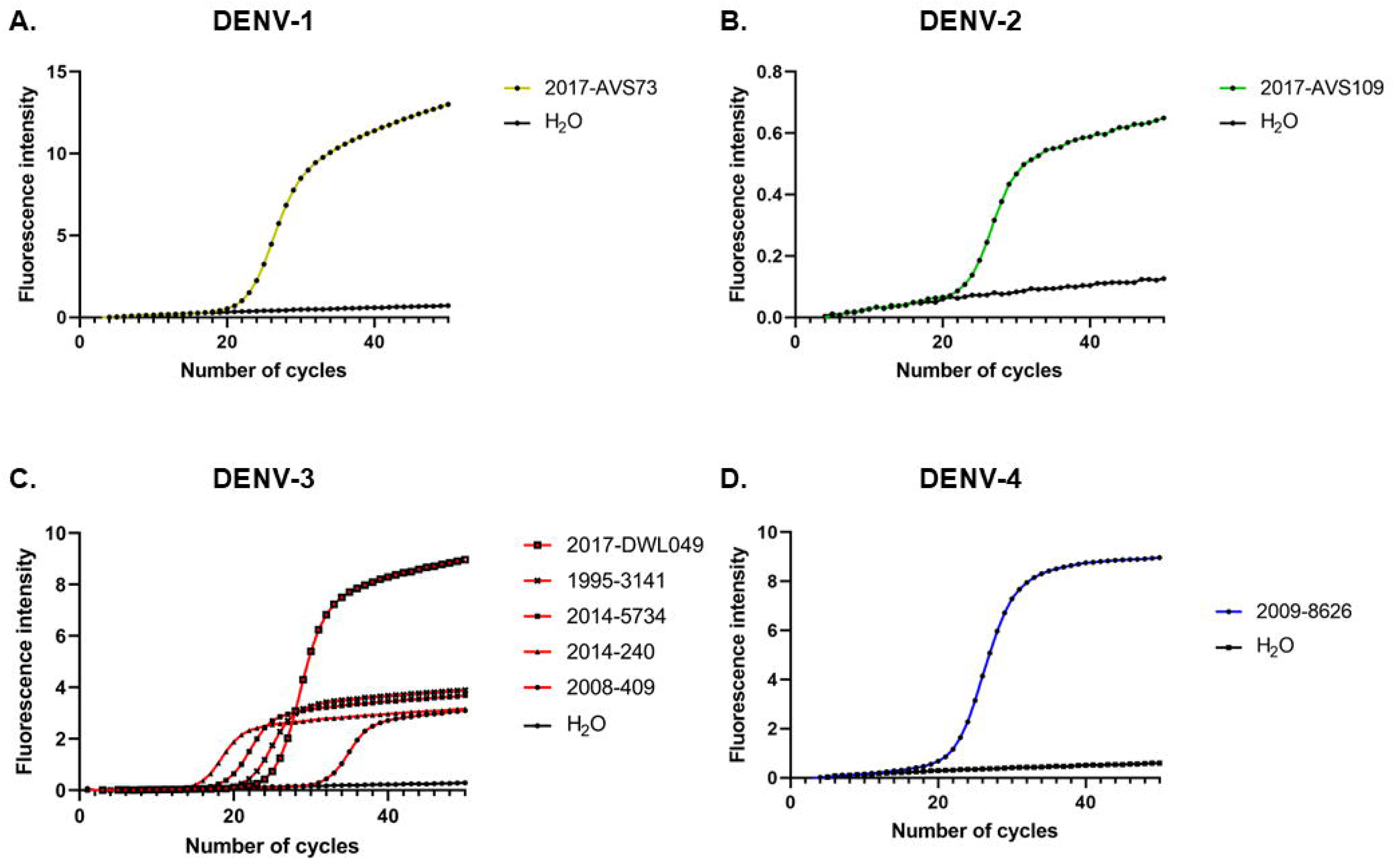
Singleplex detection of DENV-1 to -4 with optimized primers and probes. **A-D.** Fluorescence signal intensity as a function of the number of PCR cycles obtained with the optimized primers and probes are shown for DENV-1 (**A.**), DENV-2 (**B.**), DENV-3 (**C.**) and DENV-4 (**D.**).

### Multiplexing of the qRT-PCR to detect and quantify all four DENV serotypes

A color compensation experiment was implemented (Supplementary figure 2). Optimized primers and probes were then challenged in multiplex qRT-PCR. Despite color compensation, minimal crosstalk was still observed between the DENV-1, -2, -3 and -4 channels, especially in the DENV-2 channel (Figure 3A.-D.). The fluorescence threshold was thus manually adjusted in the LightCycler 480 software, based on the signal of positive controls containing a single DENV strain of each DENV-1, -2, -3 and -4 serotype. This threshold adjustment allowed to detect the strains from each serotype in the dedicated channel. The threshold was adjusted in subsequent experiments using a similar approach based on positive controls. Following color compensation and manual threshold adjustment, optimized primers and probes allowed the accurate detection of DENV-1 to DENV-4 in multiplex qRT-PCR (Figure 3E.-H.). This qRT-PCR was therefore specific, and accurately distinguished the four DENV serotypes.

**Figure 3.**
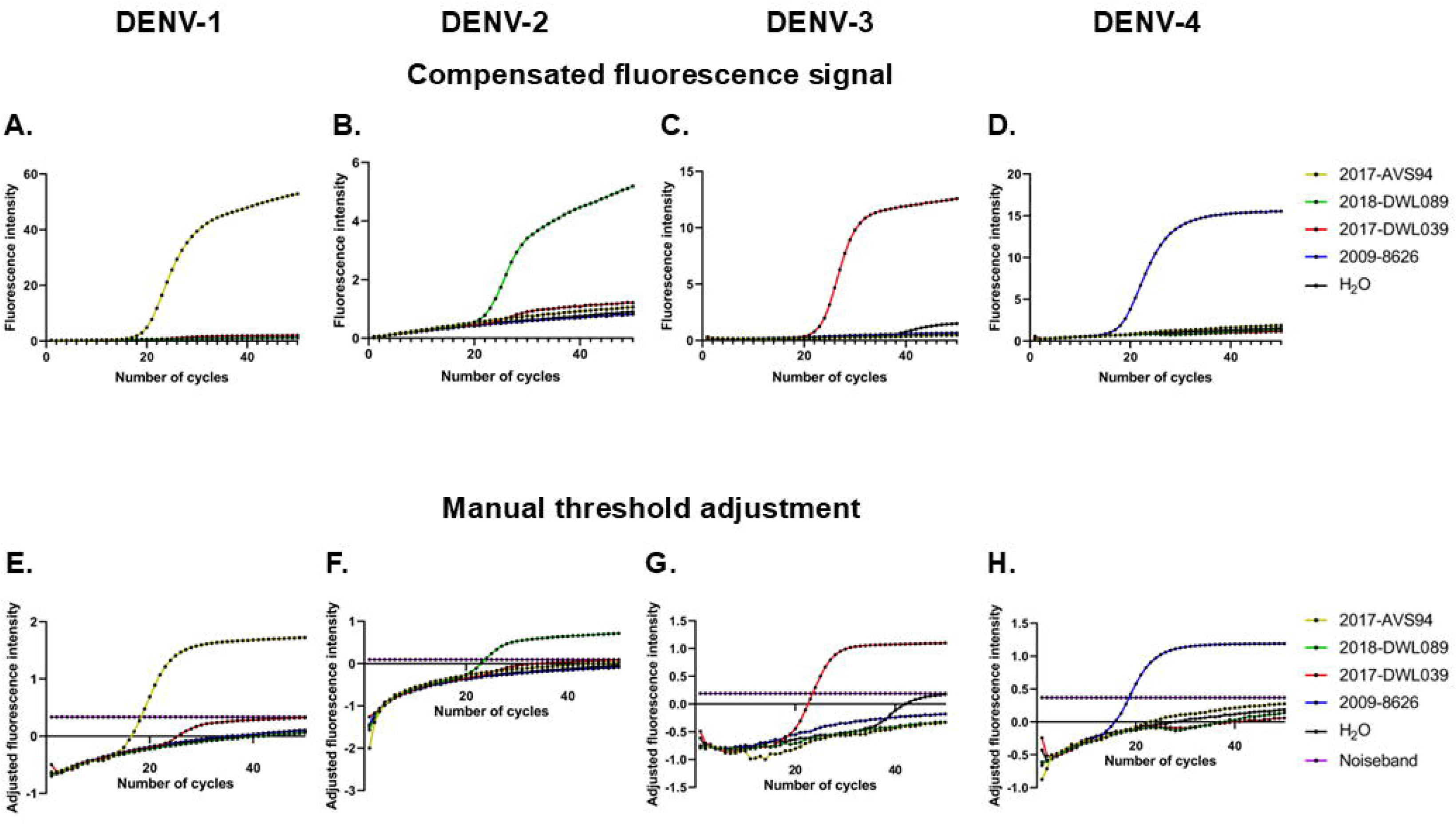
Multiplex detection of DENV-1, -2, -3 and -4 using optimized primers and probes. **A.-D.** Fluorescence signal intensity as a function of the number of PCR cycles after color compensation, for DENV-1 (**A.**), DENV-2 (**B.**), DENV-3 (**C.**) and DENV-4 (**D.**). **E.-H.** Fluorescence signal intensity as a function of the number of PCR cycles after color compensation and manual adjustment of the fluorescence signal, for DENV-1 (**E.**), DENV-2 (**F.**), DENV-3 (**G.**) and DENV-4 (**H.**).

The absolute concentration in genome copies per µL of four RNA extracts from the DENV-1 to -4 strains were determined by absolute quantification ^19^. Using these four RNA extracts of known concentration, a standard curve was established for each DENV-1 to -4 serotypes (Supplementary figure 3), allowing the absolute quantification of RNA extracts from DENV-1 to -4 with the multiplex serotype-specific qRT-PCR. This assay allowed to determine the sensitivity of this serotype-specific qRT-PCR, which was higher than 5.08×10^1^, 5.16×10^1^,7.14×10^1^ and 1.36×10^1^ genome copies/µL for DENV-1, -2, -3 and -4 respectively (Supplementary figure 3). qRT-PCR efficiency was 1.993, 1.975, 1.902, 1.898 for DENV-1, -2, -3 and -4 respectively. Linearity (R²) was 0.99975, 0.99975, 0.9985, 0.99965 for DENV-1, -2, -3 and -4 respectively.

### Quantification of DENV concentrations in a mix of two isolates belonging to different serotypes

The ability of the multiplex serotype-specific qRT-PCR to accurately quantify the concentration of two strains belonging to two different serotypes in the same RNA extract was next challenged. Concentration in viral genome copies per µL were thus determined for three viral supernatants from DENV-1, -2 and -3 serotypes. These viral supernatants were subsequently mixed two by two at ratios 0:100, 25:75, 50:50, 75:25 and 100:0. Following RNA extraction, the absolute concentration in viral genome copies per µL of each of these viral mixes was determined using the optimized quantitative multiplex serotype-specific RT-PCR. In each viral mix, the determined concentration of each strain as compared to its maximum in the 100:0 mix was not significantly different from the expected concentration (Figure 4A, *p* >0.05 in Khi² test for every determined concentration). Consistently, the determined relative percentage of each strain in the mix was not significantly different from the theoretical ratio introduced in the mix (Figure 4B, *p* >0.05 in Khi² test for every ratio).

**Figure 4.**
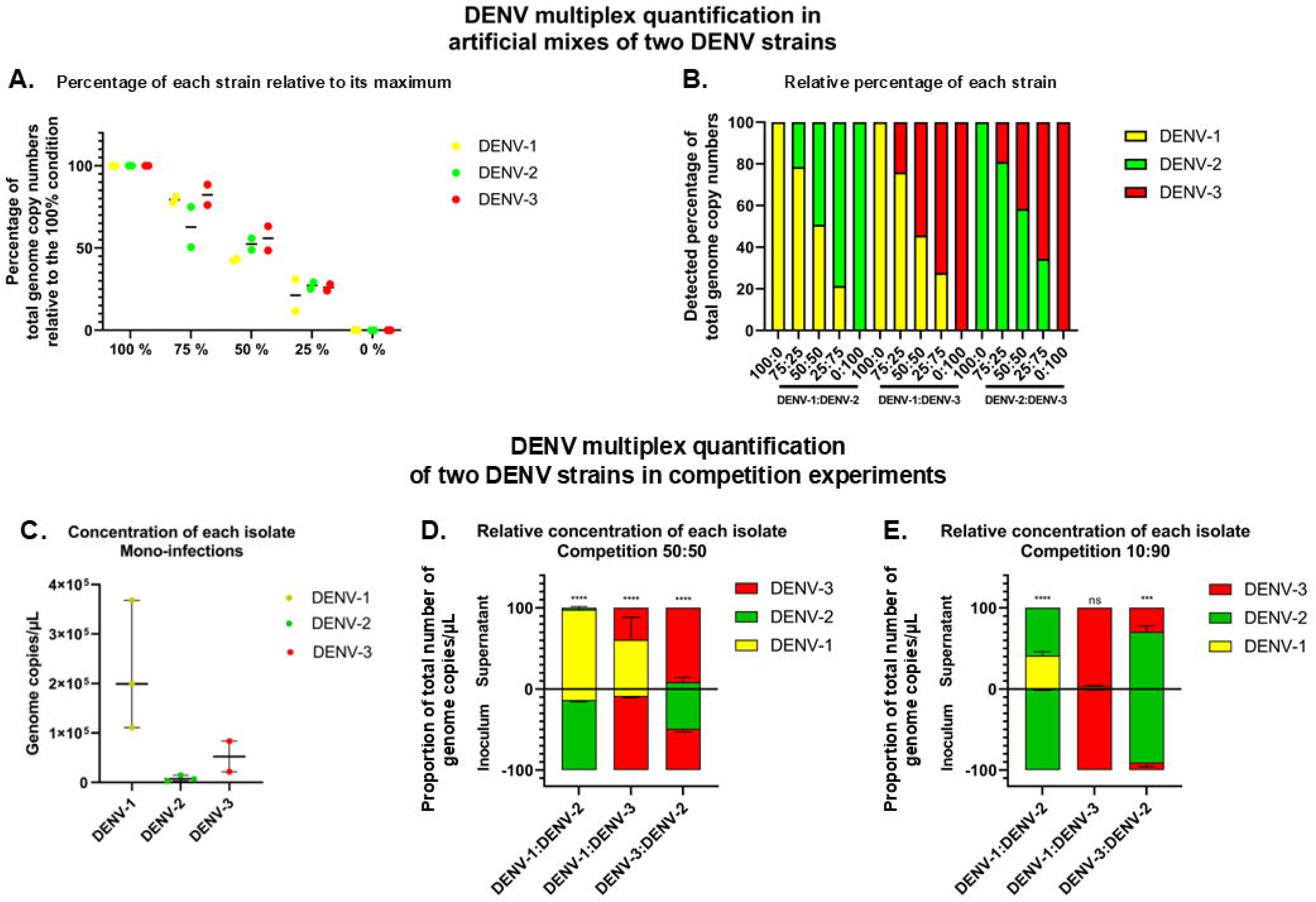
Quantification of genome copies for each DENV serotype in mixes of viral suspensions. Mixes of two viral suspensions at known ratios were subjected to RNA extraction and quantitative multiplex serotype-specific RT-PCR. **A.** Viral RNA concentrations of the three strains from DENV-1 to -3 were expressed as a percentage of viral genome copy concentrations as compared to the maximum detected for each strain at a ratio 100:0. **B.** Relative RNA concentrations of the three strains from DENV-1 to -3 were expressed as a percentage of the total number of viral genome copy concentrations detected in each mix of two strains. **C.** Absolute quantification using the quantitative serotype-specific multiplex RT-PCR of DENV-1, -2 and -3 genome copies/µL in the supernatant of HuH7 infected with a viral suspension containing each a single viral isolate of each of the three serotypes. **D-E.** Relative concentration of each isolate in the inoculum and the supernatant of HuH7 cells infected with a viral suspension containing a mix of DENV-1:DENV-2, DENV-1:DENV-3 or DENV-3:DENV-2 at a ratio 50:50 (**D.**) or 10:90 (**E.**). The concentration was determined using the quantitative serotype-specific multiplex RT-PCR. Mean and Standard Deviations of three independent triplicates are shown in **D-E**.

### Comparison of the replicative fitness of two DENV serotypes

The relative replicative fitness of three DENV strains belonging to serotype -1, -2 and -3 was estimated through mono-infections of human HuH7 cells with a viral suspension containing a single viral isolate. The quantitative multiplex serotype-specific RT-PCR showed that the DENV-1 strain exhibited the higher concentration of genome copies per µL as compared to the DENV-2 and -3 strains, suggesting a higher replicative fitness (Figure 4C).

This replicative fitness was challenged in competition assays. HuH7 cells were infected with viral suspensions containing a ratio 50:50 of a combination of two DENV strains belonging to each of DENV-1, -2 and -3 serotypes. Ratio calculation was based on the titer of viral suspension determined by immunofluorescent focus assay. The quantitative multiplex serotype-specific RT-PCR revealed that, the supernatants of DENV-1:DENV-2 and DENV-1:DENV-3 mixes contained 97.5 and 60.5 % of DENV-1 RNA copies/µL as compared to 14.8 and 9.9 % in the inoculum, respectively, evidencing a significant enrichment of the supernatant in DENV-1 RNA upon replication (Figure 4D, *p*<0.0001 in Chi² tests). Similarly, DENV-3 RNA was significantly enriched in the supernatant of cells infected with the DENV-3:DENV-2 mix (91.6% in the supernatant VS 49.7% in the inoculum, *p*<0.001 in Chi² test). This enrichment evidenced an enhanced replicative fitness of DENV-1 over DENV-2 and DENV-3, and of DENV-3 over DENV-2.

Competition assays performed with viral suspensions containing a ratio 10:90 of a combination of the same DENV strains as the one used in the 50:50 competition assay evidenced a significant enrichment in DENV-1 in the supernatant of the DENV-1:DENV-2 mix as compared to the inoculum (Figure 4E, 41.2% VS 1.3%, *p*<0.0001 in Chi² test), confirming DENV-1 enhanced replicative fitness over DENV-2. Similarly, DENV-3 was significantly enriched in the supernatant of DENV-3:DENV-2 mix as compared to the inoculum (29.8% VS 8.0%, *p<*0.001 in Chi² test), confirming DENV-3 fitness advantage over DENV-2. Only DENV-1 enhanced replicative fitness over DENV-3 was not confirm in this 10:90 competition assay.

## Discussion

In the current study, we optimized a quantitative serotype-specific RT-PCR and developed a competition assay allowing to compare the fitness of strains belonging to two different DENV serotypes using this optimized qRT-PCR. This serotype-specific qRT-PCR is easily implementable in any laboratory equipped with a real time open-system thermocycler. Quantifying specifically strains from different serotypes, genotypes or lineages can be based on Sanger sequencing, in which the height of the peak at the polymorphic position correlates with the amount of genome copies from the corresponding strain ^18^. Next Generation Sequencing approaches may also be used, in which the number of reads of each strain correlates with the number of genome copies of the corresponding strain ^22,23^. Both techniques are more expensive than a serotype-specific qRT-PCR and require both sequencing platforms and bioinformatics skills. Other qRT-PCR relying on genetic divergence between DENV strains, including between strains of different genotypes or lineages have also been developed ^24^. Of importance, this optimized multiplex serotype-specific qRT-PCR may be used in diagnostic laboratories to identify the infecting DENV serotype and could easily be deployed on the field using field-compatible real-time thermocyclers, albeit possibly requiring fluorochrome adjustments.

Competition assays allow to specifically detect significant differences in fitness between two different strains, belonging to two different serotypes for instance ^15-17^. The *in vitro* competition assay in human cells developed in the current study can now be implemented in studies of DENV serotype fitness in relevant epidemiological contexts. This competition assay design coupled to the serotype-specific qRT-PCR could be adapted to other human or mosquito cell lines. Additionally, the fitness could be followed at different timepoints over time. Finally, this serotype-specific qRT-PCR could be implemented to quantify competition assays *in vivo* in the mosquito vector.

In our competition assay, although we infected the cells with equal or 10:90 MOIs of the two viral strains, the ratio in genome copies in the inoculum was different than 50:50 or 10:90. This discrepancy is due to differences in the ratio genome copies/FFU between strains, which produce variable amounts of defective viral particles, even from one viral preparation to the next ^25^. This highlights the importance of adjusting the inoculum according to the MOI, to maintain a comparable load of infectious viruses in the inoculum. Further, performing infections with multiple ratios allows to confirm the observed trend.

This study suffers limitations. First, the accuracy of our serotype-specific quantification may be limited to the DENV lineages used to design the primers and probes. Only the sequences of DENV-3-specific primers and probes were adjusted. However, an alignment of their sequences with recent DENV-3 genomes collected in different locations (Figure 1) demonstrated that these sequences were likely adapted to detect all DENV-3 strains included in the sequence analysis. Second, the Atto590 fluorochrome used to detect and quantify DENV-2 showed a low signal. Signal-to-noise ratio could be enhanced using more efficient fluorochromes such as Alexa Fluor 594. Third, despite color compensation, we still observed some residual fluorescence cross-talk, which consequences were abrogated through a manual definition of the threshold based on the signal of positive controls. Fourth, the efficiency of our competition assay was not challenged for DENV-4. However, the study of DENV-4 was irrelevant in the recent epidemiology of dengue in New Caledonia and DENV-4 has a low circulation worldwide ^26^. The strategy described in the current study can be easily adapted to assess DENV-4 replicative fitness, should this quantification be required in a given epidemiological context. Fifth, the competition assay developed in the current study requires a Biosafety Laboratory, which may be difficult to access in less favored areas. Finally, if the fitness of several strains from each serotype was to be quantified, the experimental setup can easily become laboratory intensive.

## Conclusions

Overall, we multiplexed and optimized a serotype-specific qRT-PCR to allow the simultaneous detection and quantification of the genome copies from two viral strains belonging to two different serotypes in a viral mixture. Coupled to a relevant competition assay described in the current study, this methodology allows to compare the replicative fitness of strains belonging to two different serotypes. Such *in vitro* assessment of viral fitness may allow to evaluate the contribution of DENV replicative fitness to DENV serotype replacements in dengue-endemic or endemo-epidemic settings.

## List of abbreviations

AU: Arbitrary Units
CO_2_: carbon dioxide
°C: Celsius degrees
CA: California
DENV: dengue virus
DMEM: Dulbecco Modified Eagle Medium
FCS: Fetal Calf Serum
FFU: Focus Forming Units
H_2_O: water
mL: milliliters
MOI: multiplicity of infection
NC: New Caledonia
qPCR: quantitative Polymerase Chain Reaction
qRT-PCR: quantitative Real-Time Polymerase Chain Reaction
RNA: Ribonucleic Acid
TPB: Tryptose Phosphate Broth
UK: United Kingdom
USA: United States of America

## Declarations

### Ethics approval and consent to participate

The current study was conducted in compliance with the Declaration of Helsinki principles. Serum samples collected in 1995-2021 came from laboratory-confirmed dengue patients who did not object to the secondary use of their serum sample for research purposes. This study reusing serum samples received administrative and ethical clearance in France from the “Comité de Protection des Personnes Sud-Est II” (n° ID-RCB 2019-A03114-53, n° CPP 19.12.06.49357) and from the Consultative Ethics Committee of New Caledonia. The study was recorded on Clinicaltrials.gov (ID: NCT04615364).

### Consent for publication

Not applicable.

### Availability of data and materials

All data generated or analyzed during this study are included in this published article and its supplementary information files. Sequence data have been deposited in GenBank (MW315187, MW315194) or in the GISAID (https://gisaid.org/) EpiArbo database (EPI_SET_240904sd doi: 10.55876/gis8.240904sd).

Materials used in the current study are available from the corresponding author on reasonable request.

### Authors’ contributions

ESL, MDR and CI conceived the study and acquired funding; AFG, LR and CI performed the experiments; AFG, LR and CI analyzed the data; AFG and CI wrote the initial version of the manuscript; all authors read and approved the final manuscript.

## Acknowledgements

The authors thank Margot Petit for technical assistance.

## Supplementary materials

**Supplementary material. Sequence of the DENV RNA calibrator “iv-RNA 4”.**

**Supplementary figure 1. Standard curve for the absolute quantification of DENV RNA genome copies using an RNA calibrator.**

**A.** Compensated fluorescence intensity as a function of the number of PCR cycles for each dilution of the RNA calibrator. A signal was detected for the dilutions 10^8^ to 10^3^ copies/µL of the RNA calibrator. The limit of detection of this qRT-PCR was thus 10^3^ copies/µL. **B.** Number of PCR cycles as a function of the logarithmic transform of the concentration of the RNA calibrator. Technical duplicates were run simultaneously.

**Supplementary figure 2. Color compensations.**

**Supplementary figure 3. Standard curves.**

**Supplementary figure 4. Singleplex qRT-PCR for the detection of DENV-1 to -4 with the initial primers and probes. A-D.** Fluorescence signal intensity as a function of the number of PCR cycles obtained with the initial primers and probes are shown for DENV-1 (**A.**), DENV-2 (**B.**), DENV-3 (**C.**) and DENV-4 (**D.**).

