## Supplementary figures and images for "Development of a competition assay to assess the *in vitro* fitness of dengue virus serotypes using an optimized serotype-specific qRT-PCR"

### Supplementary Figure 1

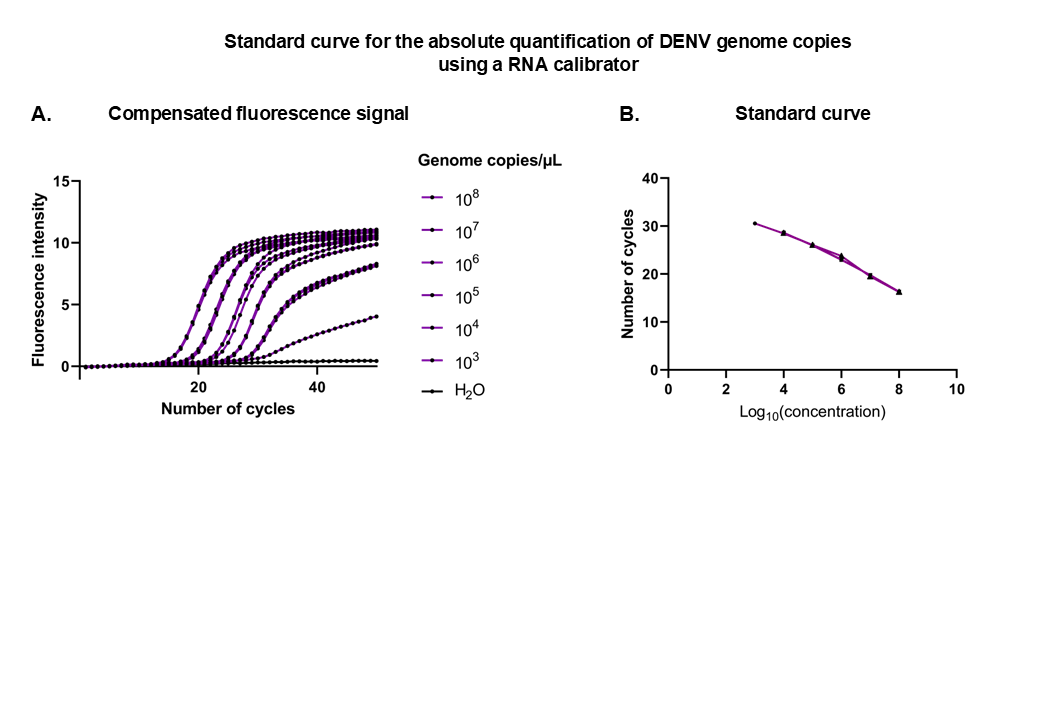

### Supplementary Figure 2

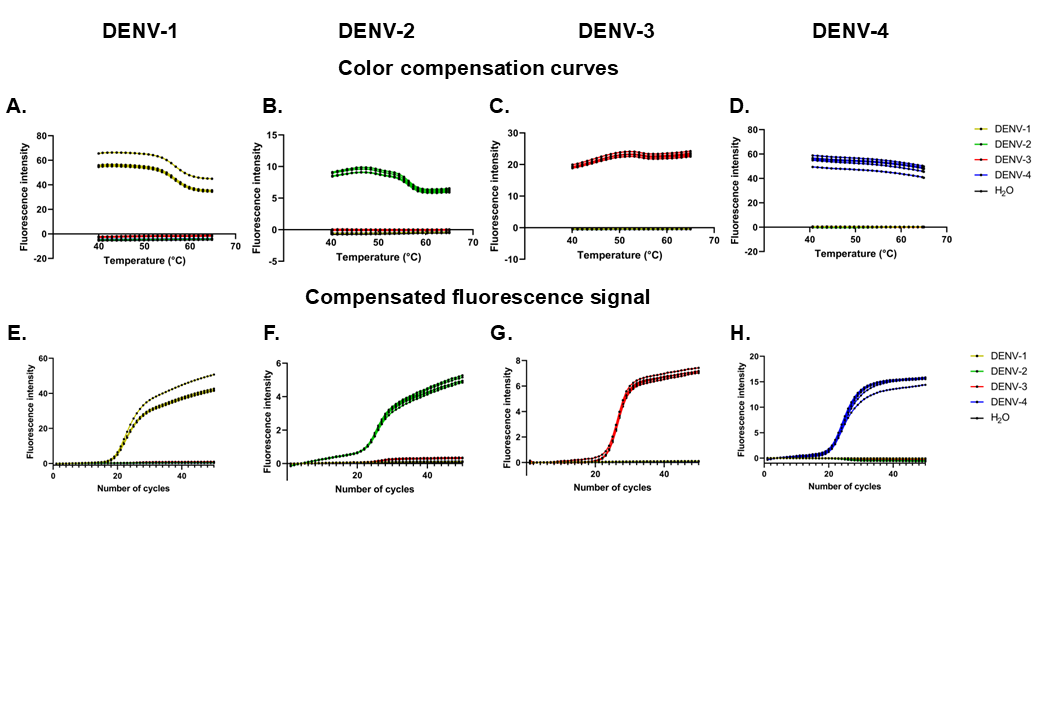

### Supplementary Figure 3

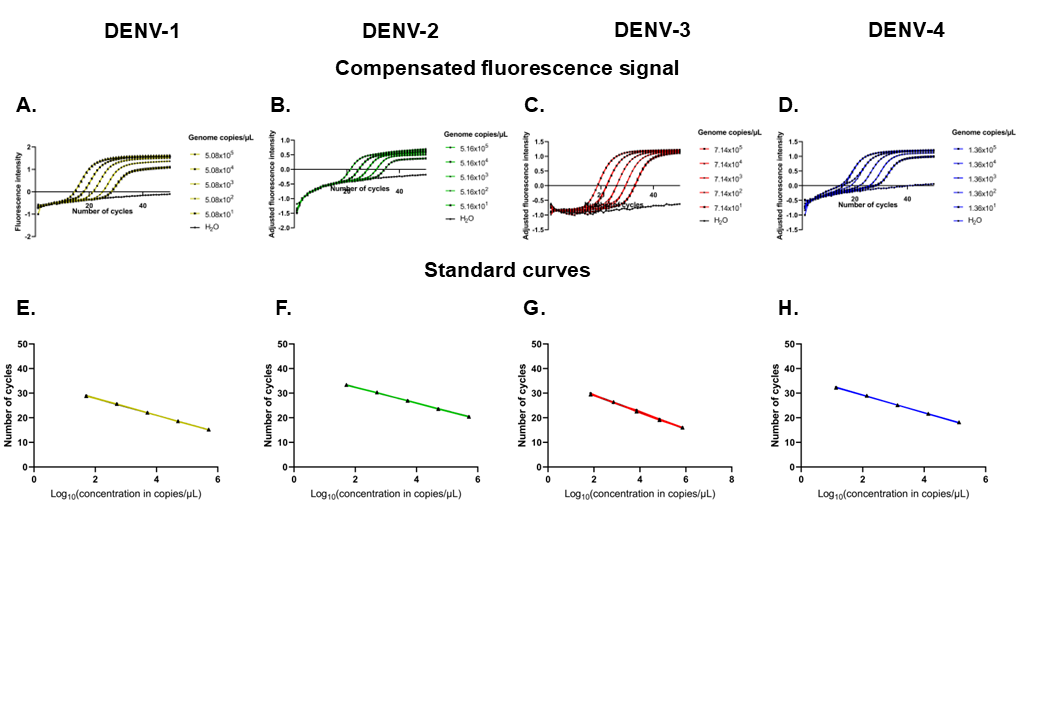

### Supplementary Figure 4

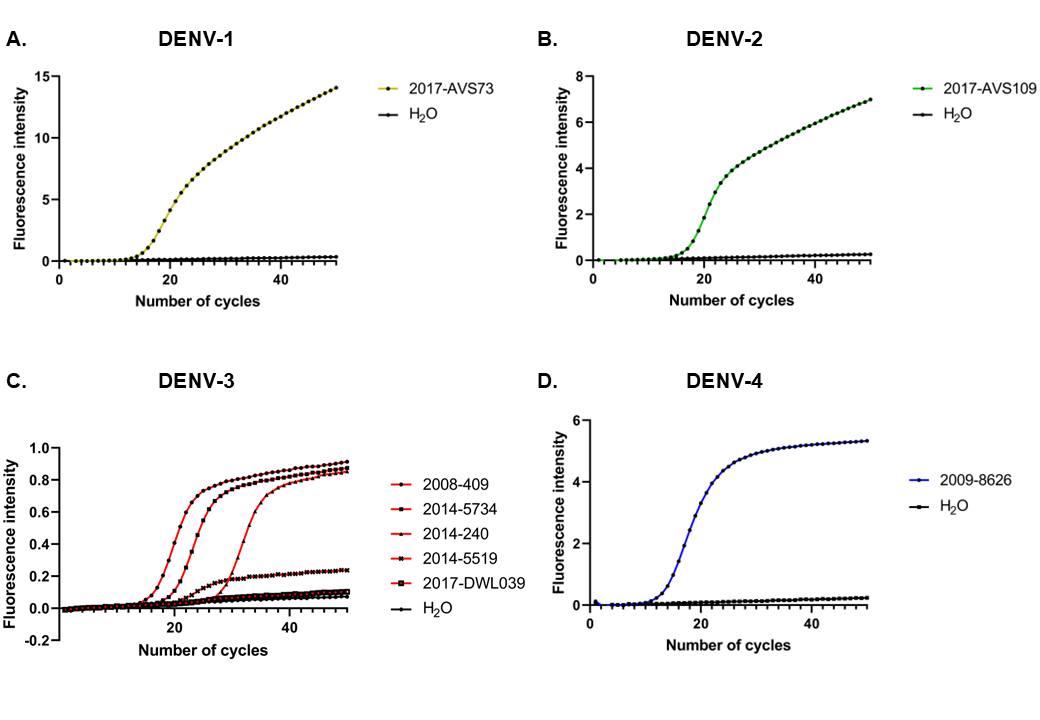
