## Supplementary material for "Development of a competition assay to assess the *in vitro* fitness of dengue virus serotypes using an optimized serotype-specific qRT-PCR"

**Sequence of the DENV RNA calibrator “iv-RNA 4”:**

gaattcgcccttatccatgcccatcaccaatggatgacaacagaagacatgttgtcagtgtggaatagggtttggatagaggaaaacccatggatggaggataaaacccatatatccagttgggaagatgttccatacttaggaaaaagggaagatcagtggtgtggatccctgataggcttaacagcaagggccacctgggccactaatatacaagtggccataaaccaagtgagaaggcttattgggaatgagaattatctagattacatgacatcaatgaagagattcaagaatgagagtgatcccgaaggggcactctggtaagtcaacacattcacaaaacaaaggaaaataagaaatcaaacaaggcaagaagtcaggccggattaagccatagtacggtaagagctatgctgcctgtgagccccgtctaaggacgtaaaatgaagtcaggccggaagccacggtttgagcaaaccgtgctgcctgtagctccatcgtggggatgtaaaaacccgggaggctgcaacccatggaagctgtacgcatggggtagcagactagtggttagaggagacccctcccaaaacacaacgcagcagcggggcccaacaccaggggaagctgtaccctggtggtaaggactagaggttagaggagaccccccgcataacaataaacagcatattgacgctgggagagaccagagatcctgctgtctctacagcatcattccaggcacagaacgccagaaaatggaatgaagggcgaattc
